# The STAT3-VDAC1 Axis Modulates Mitochondrial Function and Plays a Critical Role in the Survival of Acute Myeloid Leukemia Cells

**DOI:** 10.1101/2024.06.24.600435

**Authors:** Kellen B. Gil, Jamie Borg, Rosana Moreira Pereira, Anagha Inguva-Sheth, Geovana Araujo, Jeremy Rahkola, William Showers, Abby Grier, Angelo D’Alessandro, Clayton Smith, Christine McMahon, Daniel A. Pollyea, Austin E. Gillen, Maria L. Amaya

**Author notes:** Corresponding Author’s Contact: University of Colorado. Rocky Mountain Regional VA Medical Center. 1700 N. Wheeling St. Building B. Aurora CO 80045. Competing Interests Statement The authors declare no competing financial interests.

## Abstract

Signal transducer and activator of transcription 3 (STAT3) is a well-described transcription factor that mediates oxidative phosphorylation and glutamine uptake in bulk acute myeloid leukemia (AML) cells and leukemic stem cells (LSCs). STAT3 has also been shown to translocate to the mitochondria in AML cells, particularly when phosphorylated at the serine 727 (pSTAT3 S727) residue. Inhibition of STAT3 results in impaired mitochondrial function and decreased leukemia cell viability. We discovered a novel interaction of STAT3 with voltage-dependent anion channel 1 (VDAC1) in the mitochondria that provides a mechanism through which STAT3 modulates mitochondrial function and cell survival. Through VDAC1, STAT3 regulates calcium and oxidative phosphorylation in the mitochondria. STAT3 and VDAC1 inhibition also results in significantly reduced engraftment potential of LSCs, including primary samples resistant to venetoclax. These results implicate STAT3 as a therapeutic target in AML.

## Introduction

Acute myeloid leukemia (AML) is a genetically heterogenous and highly aggressive myeloid neoplasm with poor prognosis^1,2^. Standard therapy for AML has historically consisted of induction chemotherapy with an anthracycline and cytarabine, followed by consolidation with either hematopoietic stem cell transplant or high dose cytarabine^3^. Recently, therapeutic options have broadened with the advent of novel targeted therapies^4–7^. However, despite high response rates, relapse is common^6^. Relapsed disease is believed to originate from a quiescent subpopulation of therapy-resistant leukemic stem cells (LSCs)^8^ which are found in greater abundance at the time of relapse compared to diagnosis^9–12^, and negatively correlate with survival^10,11^. LSCs demonstrate a unique vulnerability in their preferential reliance on mitochondrial activity and oxidation phosphorylation (OXPHOS)^12–14^. While Bcl-2 inhibition with venetoclax (Ven) in combination with the hypomethylating agent (HMA) azacitidine has demonstrated selectivity for LSCs via inhibition of OXPHOS^13^, resistance frequently develops via alterations in mitochondrial metabolism or activation of alternative antiapoptotic pathways^15–19^. Further, prior studies of patients who progress after frontline HMA/Ven have shown very poor outcomes, with a median survival following failure of HMA/Ven of 3 months or less^20,21^. New strategies targeting LSCs via their reliance on OXPHOS are of significant interest and have been described in several reports^7,13,22^, however further research is needed to elucidate the mechanisms underlying these observations.

Signal transducer and activator of transcription 3 (STAT3) has been shown to be important for leukemogenesis and is known to be highly expressed in many AML patient samples and cell lines^23–26^. Canonically, STAT3 is known to undergo phosphorylation at residue Tyr^705^ leading to dimerization and translocation to the nucleus where it functions as a transcription factor regulating cell development, renewal, proliferation, and cell death^24,27–29^. Our previous work additionally established that STAT3’s transcriptional activity regulates mitochondrial function via a *MYC*-SLC1A5-mediated pathway^26^. Despite its well-described nuclear role as a transcription factor, STAT3 has also been discovered to localize to the mitochondria^30,31^. Prior work has suggested a variety of functions in the mitochondria, including modulation of electron transport chain (ETC) activity^30–32^, regulation of mitochondrial genes^33^, and regulation of mitochondrial calcium flux^34,35^. While phosphorylation of STAT3 at both Tyr^705^ (pSTAT3 Y705) and Ser^727^ (pSTAT3 S727) sites have been found in the mitochondria^30–32,35,36^, Ser^727^ phosphorylation is critical for modulation of mitochondrial functions such as ETC activities^30,31^. These data suggest that STAT3 plays a critical role in the mitochondria, although this role in AML is not well characterized.

Here we show STAT3 plays a key role in mitochondrial function of AML cells, and that it interacts with mitochondrial proteins including voltage-dependent anion channel 1 (VDAC1), playing a regulatory role of mitochondrial calcium and OXPHOS.

## Materials & Methods

### Cell Culture

For human primary AML samples, base media of minimum essential medium (StemCell Technologies 09655) and 5.5 mM of glucose was used, supplemented with physiologic levels of amino acids as previously reported^12,26^ and 10 nM of human cytokines stem cell factor (SCF) (Peprotech 300-07-100), interleukin-3 (Peprotech 200-03-50), and Fms Related Receptor Tyrosine Kinase 3 (FLT3) (Peprotech 300-19-100). The MOLM-13 cell line was used, which was purchased from the University of Colorado Cell Technologies Shared Resource (CTSR), which was obtained from DSMZ.

### Patient Samples and LSC Enrichment

Primary human AML samples were obtained from apheresis products or bone marrow of patients with AML who gave written consent for sample procurement at the University of Colorado, according to the Colorado IRB Protocol #12-0173. LSCs were isolated using a ROS-low strategy^37^. For analysis of LSC-enriched fractions, specimens were processed as previously described^12,38^.

### Viability Measured by Flow Cytometry

Cell viability was measured using Annexin V (BD 556421) and Dapi (BD 564907) stains and measured by flow cytometry. Ghost dye (Tonbo Bioscience 130865T100) was also used to confirm cell death and measured by flow cytometry.

### Single Cell RNA Sequencing

Clusters were annotated using clustifyr 1.9.1^39^ and the leukemic/normal bone marrow reference dataset generated by Triana and colleagues^40^. Scanpy and Seurat 4.1.1 were then used to generate uniform manifold approximation and projections from the TotalVI embeddings and perform exploratory analysis, data visualization, etc^41^.

### Flow Cytometry ImageStream

FIX & PERM cell fixation assay (ThermoFisher GAS004) was used to stain for intracellular markers pSTAT3 S727 (BD 565416) and Tomm20 (Abcam ab205486). Cells were then washed with FACS Buffer (1% FBS in 1X PBS) and ran on the ImageStreamX MkII (Cytek Biosciences). Samples were then analyzed using the Ideas 6.2 software threshold masking for mitochondrial staining which was utilized to determine the intensity per area of pSTAT3.

### Mitochondrial Oxygen Flux Analysis

Oxygen consumption rate (OCR) was measured using the Seahorse XF96 Cell Mito Stress Test kit (Agilent 103015-100) on the Seahorse XFe96 Extracellular Flux Analyzer (Agilent) according to manufacturer’s protocol. Treated MOLM-13 cells were washed and plated on Cell-Tak (Corning 354240) coated XFe96 cell culture microplates (Agilent) at 150,000 cells/well in 5 replicates using Seahorse Assay RPMI Medium (Agilent). OCR was measured at basal level and after injection of 5 μg/mL oligomycin, 2 μM Carbonyl cyanide-4-(trifluoromethoxy)phenylhydrazone (FCCP), 5 μM antimycin A, and 5 μM rotenone.

### Metabolomic Experiments

Metabolomics analyses were performed on 10 μl of sample extracts via ultra-high pressure liquid chromatography coupled to high resolution mass spectrometry (Vanquish – QExactive – Thermo Fisher, San Jose, CA, USA) using a high-throughput 5 min gradient-based method.

### siRNA Transfections

siRNA sequences targeting STAT3 (siSTAT3) and a scrambled control (siSCR) were purchased directly from Horizon Discovery’s ON-TARGETplus siRNA Reagents collection (L-003544-00-0005 (Stat3) and D-001810-10-05 (SCR)). The lyophilized siRNA products were resuspended in RNAse-free water at 5 μmol/L, which was used as a stock solution. 2×10^6^ cells were suspended in 80 μL of Buffer T, and 20μL of siRNA stock solution was added. These cells were then electroporated using the Neon Electroporation Transfection System (Thermo) according to the manufacturer’s protocol using the following settings: 1,600 V, 10 ms, 3 pulses.

### Electron Microscopy

MOLM-13 cells were treated with DMSO or Stattic 5μM (Sigma-Aldrich S7947) for 12 hours and subsequently fixed prior to submission to the Electron Microscopy Core at University of Colorado for subsequent imaging. Mitochondrial number and area were measured using the ImageJ software.

### Immunoprecipitation Assay

Mitochondrial protein was isolated from cells (Thermo 89874) and immunoprecipitation assays were performed using the mitochondrial protein IP kit from Abcam (catalog # ab239710) according to the manufacturer’s protocol. Digitonin was used as the detergent (Sigma D141-100). Solubilized mitochondrial supernatants were incubated overnight at 4°C in 2ug primary antibody (Stat3 (Invitrogen MA1-13042), VDAC1/Porin (Proteintech 55259-1-AP), Mouse IgG (Millipore pp54), Rabbit IgG (Millipore PP64B). Proteins were bound using Protein G Mag Sepharose Beads (Cytiva 28944008) and eluted using RIPA buffer (Sigma-Aldrich R0278). Eluent was submitted to the University of Colorado proteomics core.

### Mitochondrial Calcium Assay

Cells were incubated with Rhod2AM (Invitrogen, R1245MP; 500nM) for 30 minutes at 37°C. Cells were then washed with calcium free, magnesium free 1X PBS (Corning, 21-031-CV) twice and resuspended in 1X calcium free, magnesium free PBS supplemented with 2% FBS (Atlas Biologicals, no. F-500-D) and analyzed by flow cytometry (PE channel). Detection for Rhod-2 AM was done on 552/581 channel. Viability Measured by Flow Cytometry Cell viability was measured using Dapi and measured by flow cytometry. Cells were stained in PBS supplemented with 2% FBS.

### Mouse Studies

Human primary AML cells were treated with either vehicle control, 5μM of Stattic or 400μM of DIDS overnight. On the same day, NSG-SGM3 mice (Jackson Laboratory 013062) were conditioned with 25 mg/kg busulfan (Alfa Aesar J61348) via intraperitoneal injection. On the second day, AML cells were washed with FACS buffer and resuspended in saline with the addition of anti-human CD3 antibody (OKT3 BioXCell BE0001-2) at a final concentration of 1 μg/10^6^ cells and incubated for 15 minutes prior to injection as a means to reduce graft versus host disease. 8-9 mice per group were injected with 2.5 x 10^6^ cells/mouse. Mice were sacrificed after 8-12 weeks and femurs were collected and flushed with FACS buffer. Flow cytometry was then performed after staining with mouse (BD 560510) and human (BD 561865) specific CD45 antibodies. All animal experiments were approved by the Rocky Mountain Regional VA Medical Center (Institutional Animal Care and Use Committee) under protocol number CD2114M.

### Western Blots

Cells were lysed in RIPA Buffer (Sigma-Aldrich R0278) supplemented with HALT Protease and Phosphatase Inhibitor Cocktail (Thermo 78442). Proteins were separated by SDS-PAGE gel (Bio-Rad 4561094), transferred to PVDF membrane (Millipore 03010040001), and blocked for 1 hour at room temperature using 5% w/v BSA in 1% Tween-20-TBS Buffer. Membranes were incubated overnight at 4°C with primary antibody against pStat3 Ser727 (CellSignaling 9134S), Stat 3 (Cell Signaling 12640S), VDAC1/Porin (Abcam ab14734), CoxIV (CellSignaling 4850S) or β-Actin (SantaCruz Biotechnology SC-47778). Membranes were washed and incubated with 1:10,000 secondary antibody against mouse (Abcam ab205719) or rabbit (Abcam ab205718) for 1 hour at room temperature. The band were visualized using LumiGlo Chemiluminescent Substrate System (SeraCare 5430-0040) and the ChemiDoc MP Imaging System (Bio-Rad).

## Data Availability

Data collected in this study were generated by authors and are available upon request.

## Results

### STAT3 localizes to the mitochondria, and it interacts with VDAC1

To investigate the role of STAT3 in human AML cells, we utilized both primary AML samples donated by patients at the University of Colorado and the AML cell line MOLM-13. Our prior work has demonstrated significantly higher STAT3 expression and phosphorylation in AML patient samples compared to normal cord blood mononuclear cells^26^. Further, high STAT3 expression appears to be enriched in leukemia stem cells from patient samples resistant to venetoclax and azacitidine based on scRNA sequencing (**Supplemental Figure 1A**), suggesting a possible role at diagnosis but also at the time of relapse. To understand the role of STAT3 in the mitochondria of AML cells, we first assessed how frequently STAT3 localizes to the mitochondria of AML cells. We utilized primary patient samples and performed ImageStream. Using an antibody specific for phosphorylated STAT3 at S727, we were able to quantify its localization to the mitochondria of primary AML samples, which occurs in over 80% of the cells as shown in **Figure 1A**. Given phosphorylation at S727 has been shown to be the critical activation site for mitochondrial STAT3, we also assessed whether isolated leukemia stem cells had increased phosphorylation at that site. pSTAT3 expression was higher in LSCs isolated from 4 primary AML compared to other bulk leukemia cells (**Figure 1B**). To further investigate the role of STAT3 in the mitochondria, we used the cell line MOLM-13. These cells also show high expression of STAT3, pSTAT3 and localization of STAT3 to the mitochondria (**Supplemental Figures 1B-D**). To identify whether STAT3 interacts with other mitochondrial proteins, we first performed immunoprecipitation assays in mitochondrial fractions of MOLM-13 cells with a STAT3 antibody, followed by mass spectrometry (**Figure 1C**). With this assay, we identified 6 mitochondrial proteins that interact with STAT3 in AML cells (**Figure 1D**), including Voltage-dependent anion-selective channel 1 (VDAC-1) (**Figure 1E**). VDAC1 is an outer mitochondrial membrane (OMM) protein which is known to play physiologic roles in regulating OXPHOS^42^ and apoptosis^43^. No prior evidence has demonstrated that mitochondrial STAT3 and VDAC1 interact, we therefore sought to further verify this novel interaction. To further confirm this interaction, co-immunoprecipitation assays from mitochondrial fraction isolates were performed using either STAT3 or VDAC1 antibodies, followed by western blot analysis. As shown in **Figure 1F**, STAT3 pull down (left) showed prominent signal for both STAT3 and VDAC1 proteins relative to IgG control, and VDAC1 pull down (right) similarly showed prominent VDAC1 and STAT3 proteins compared to IgG control by western blot.

**Figure 1.**
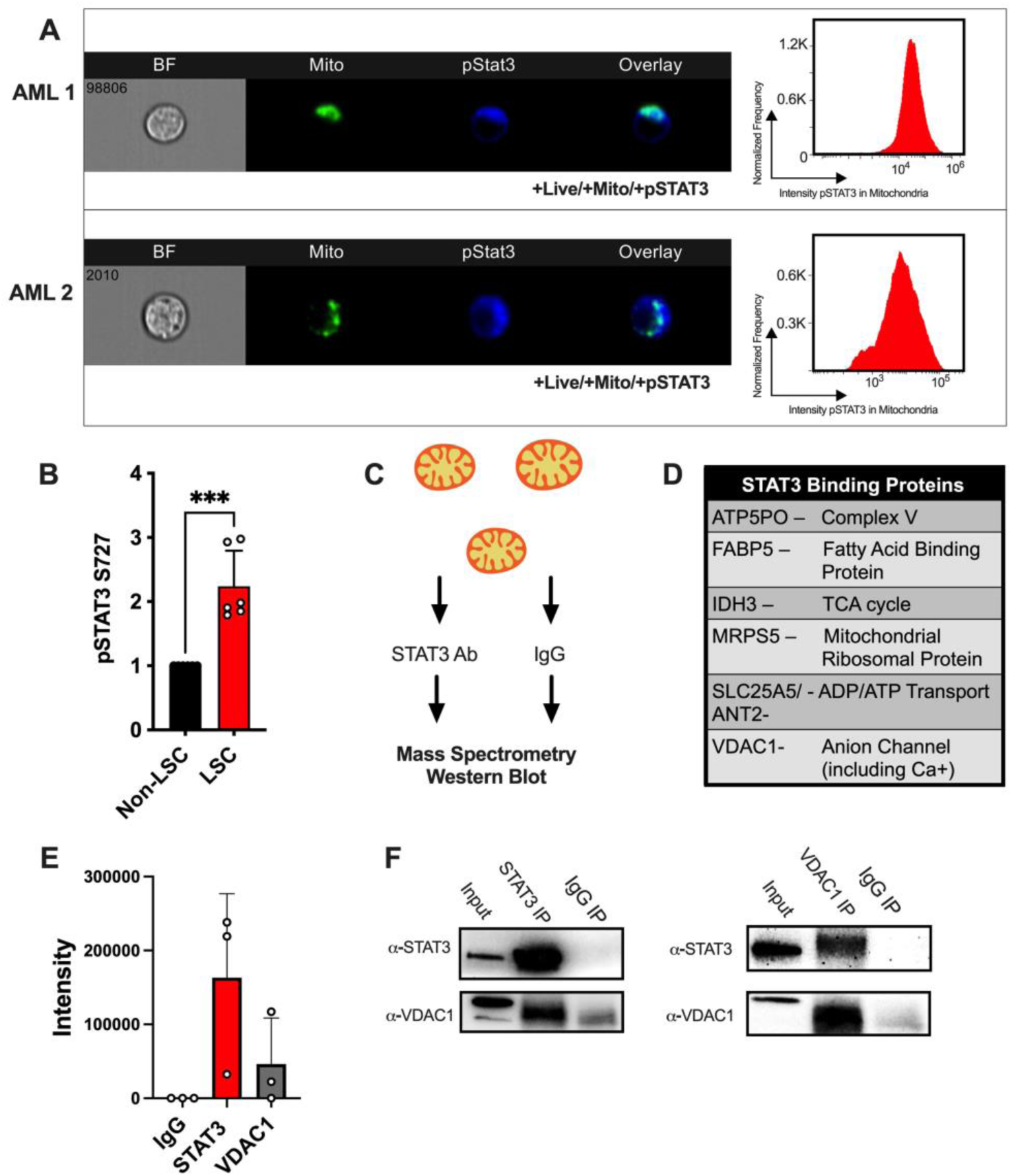
STAT3 localizes to the mitochondria and it interacts with mitochondrial proteins. (A) Representative ImageStream from two AML patient samples staining for mitochondria, pSTAT3 S727, and their respective overlays. In AML 1, 87.5% of cells were positive for pSTAT3 S727 in the mitochondria out 50,205 events. In AML 2, 81.7% of the cells were positive for pSTAT3 S727 in the mitochondria out of 41,615 events. (B) Western blot quantification of pSTAT3 at S727 comparing leukemia stem cells to bulk cells of 4 primary AML samples. (C) Cartoon representation of the experimental design for immunoprecipitation experiments. (D) Table outlining proteins interacting with STAT3 in the mitochondria of MOLM-13 cells based on mass spectrometry (E) Intensity based absolute quantification of VDAC1 protein bound to STAT3. IgG serves as a negative control and STAT3 as a positive control. (F) Western blots from STAT3 pulldown (left) showing STAT3 and VDAC1 protein expression, and VDAC1 pulldown (right) showing STAT3 and VDAC1 protein expression. Statistical analyses were performed using a Student’s t-test. P values are represented as follows: * p ≤ 0.05, ** p ≤ 0.01, *** p ≤ 0.001.

### STAT3 inhibition results in decreased mitochondrial VDAC1 resulting in calcium imbalance

To help determine the relationship between STAT3 and VDAC1 in the mitochondria, we next sought to study the effects of STAT3 inhibition. To inhibit mitochondrial STAT3, we used a potent STAT3 inhibitor, Stattic. Stattic has been shown to inhibit STAT3 dimerization, but it has also been shown to inhibit phosphorylation at S727 at higher doses, which is critical for STAT3 mitochondrial localization and function^44^. To find the optimal dose to study STAT3 inhibition in these cells, we first treated MOLM-13 cells with increasing doses of Stattic. As shown in **Supplemental Figure 2A**, 5μM of Stattic results in significant cell death at 24 hours compared to vehicle control. However, this dose did not cause significant cell death until 16 hours (**Supplemental Figure 2B**), allowing a window to understand the effects of STAT3 inhibition prior to cell death. We next tested the effects of Stattic inhibition on STAT3 in MOLM-13 cells. As shown in **Figure 2A**, Stattic treatment resulted in significant reduction of pSTAT3 S727 after 9 hours in culture, while only resulting in a limited reduction of total STAT3 (**Supplemental Figure 2C**). Since phosphorylation of S727 has been linked to localization and activity of mitochondrial STAT3, we further confirmed that culture with Stattic resulted in decreased localization of pSTAT3 to the mitochondria (**Figure 2B**). Further, western blot analysis revealed that Stattic treatment resulted in decreased levels of VDAC1 in mitochondrial fraction isolates at 9 hours, while whole cell VDAC1 had no significant changes (**Figure 2C and Supplemental Figure 2D**), suggesting that STAT3 inhibition results in decreased VDAC1 mitochondrial localization. Similar results were seen upon STAT3 knockdown by siRNA, where decreased STAT3 expression resulted in decreased mitochondrial VDAC1 (**Supplemental Figure 2E-G**). Prior work has demonstrated that both STAT3 and VDAC1 play roles in mitochondrial calcium regulation^34,35,45,46^. Additionally, mitochondrial calcium homeostasis has been shown to be important for LSCs in AML^19^. To determine the potential effects of STAT3 inhibition on mitochondrial calcium flux in AML, we performed mitochondrial calcium assays with Rhod-2 AM staining. We found that pharmacologic (Stattic) and genetic inhibition of STAT3 in MOLM-13 cells resulted in significantly decreased mitochondrial calcium levels (**Figure 2D-E** and **Supplemental Figure 2H**). Direct inhibition of VDAC1 with two inhibitors, 4,4’-diisothiocyanostilbene-2,2’-disulfonic acid (DIDS) (**Figure 2F**), and NSC (**Supplemental Figure 2I**) also resulted in decreased mitochondrial calcium compared to vehicle control. To understand whether STAT3’s regulation of mitochondrial calcium is through VDAC1, we overexpressed VDAC1 in MOLM-13 cells and treated them with Stattic to inhibit STAT3. As shown in **Supplemental Figure 2J**, a transient overexpression of VDAC1 results in a 1.5 fold increase in protein expression in MOLM-13 cells. This higher expression is enough to partially rescue the mitochondrial calcium levels upon STAT3 inhibition (**Figure 2G**).

**Figure 2.**
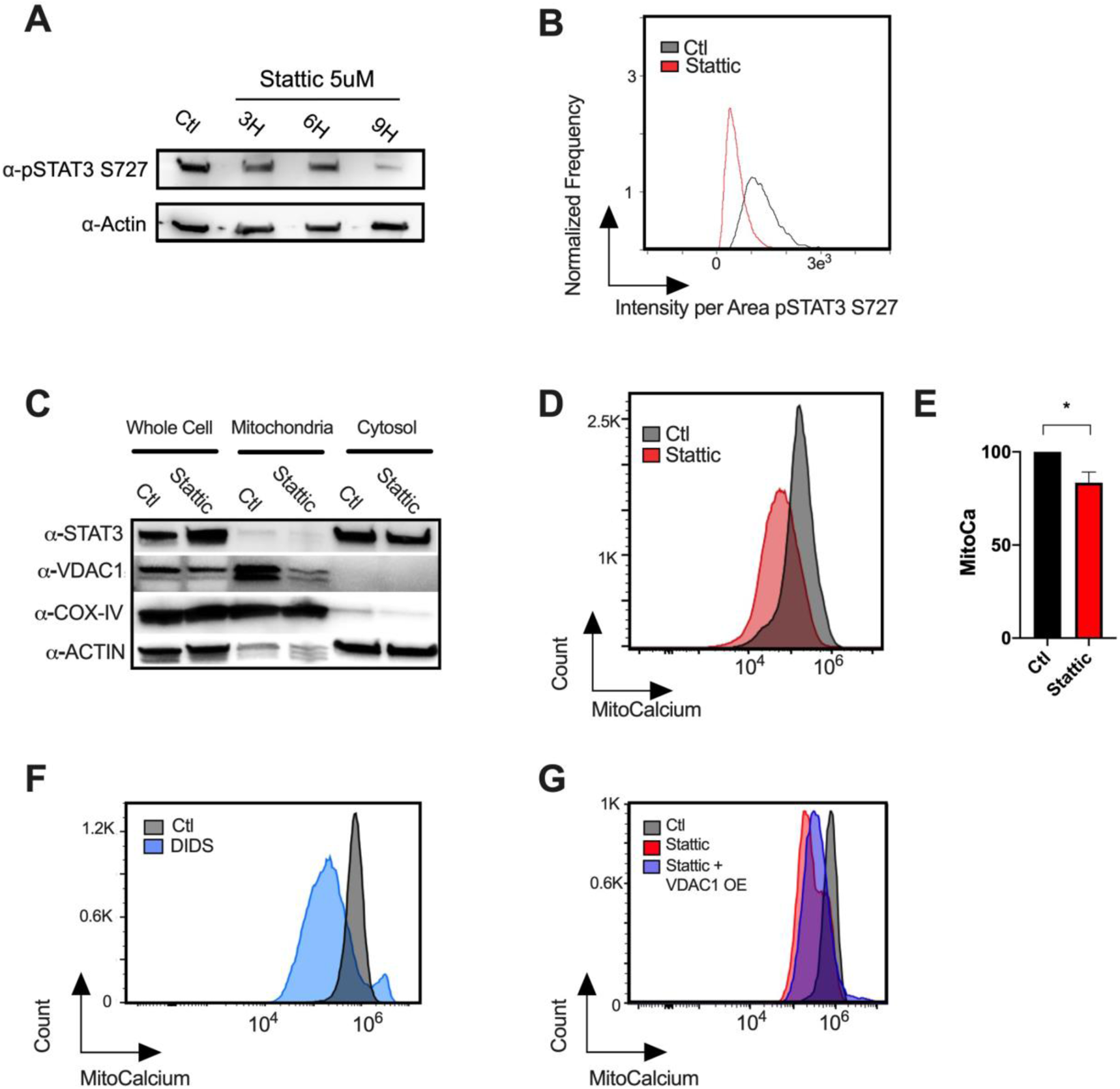
STAT3 inhibition results in decreased mitochondrial VDAC1. (A) Western blot showing pSTAT3 S727 in MOLM-13 cells in the presence of Stattic 5μM at 3, 6 or 9 hours compared to vehicle control. (B) Intensity per area of pSTAT3 S727 in the mitochondria of MOLM-13 cells treated Stattic or vehicle control as measured by ImageStream flow cytometry. (C) Western blot showing protein expression of STAT3 and VDAC1 in whole cell, mitochondrial or cytosolic fractions of MOLM-13 cells treated with Stattic 5μM or vehicle control for 9 hours. COX-IV and Actin antibodies serve as mitochondrial and cytosolic controls, respectively. (D) Mitochondrial Calcium as measured by flow cytometry in MOLM-13 cells treated with Stattic 5μM or vehicle control for 9 hours. (E) Quantification of 3 technical replicates of mitochondrial calcium in MOLM-13 cells treated with Stattic 5μM or vehicle control for 9 hours. (F) Mitochondrial calcium as measured by flow cytometry in MOLM-13 cells treated with DIDS 400μM or vehicle control for 9 hours (G) Mitochondrial calcium as measured by flow cytometry in sham-electroporated vs. VDAC1 overexpressing plasmid electroporated Molm-13 cells treated with Stattic 5μM or vehicle control for 9 hours. Statistical analyses were performed using a Student’s t-test. P values are represented as follows: * p ≤ 0.05, ** p ≤ 0.01.

Together, these studies suggest that STAT3 binds to and regulates the function of VDAC1, and that some of mitochondrial STAT3’s downstream effects may be mediated through VDAC1.

### STAT3 and VDAC1 inhibition results in reduction of OXPHOS and mitochondrial membrane potential

To assess the effects of pharmacologic inhibition of STAT3 on mitochondrial function, we first used Seahorse MitoStress testing to measure oxygen consumption rates (OCR) in the presence or absence of Stattic or DIDS. Both Stattic and DIDS treatment resulted in a significant decrease in OCR after 6 hours compared to control samples (**Figure 3A**). Consistent with this finding, inhibition of STAT3 or VDAC1 results in a mild decrease in mitochondrial reactive oxygen species (**Figure 3B-C**), suggesting less ROS is being produced due to lower TCA cycle and/or electron transport chain activity function. Metabolomics analysis of MOLM-13 samples treated with Stattic or DIDS showed multiple metabolic pathways are affected (**Supplemental Figure 3A-D**), and both resulted in abundant glutathione (**Figure 3D)** and a normal glutathione to glutathione disulfide ratio (**Figure 3E-F**), consistent low oxidative stress. These effects are likely related to decreased mitochondrial calcium, which is critical for the function of several TCA cycle enzymes^47^. Given mitochondrial calcium is also involved in regulating the mitochondrial membrane potential independently of OXPHOS^35^, we assessed this by TRME stains upon treatment with Stattic of DIDS. As shown in **Figure 3G**, inhibition of STAT3 and VDAC1 result in decreased mitochondrial membrane potential. These findings suggest that STAT3 and VDAC1 are involved in regulating TCA cycle activity while also affecting the mitochondrial membrane potential, likely through calcium regulation.

**Figure 3.**
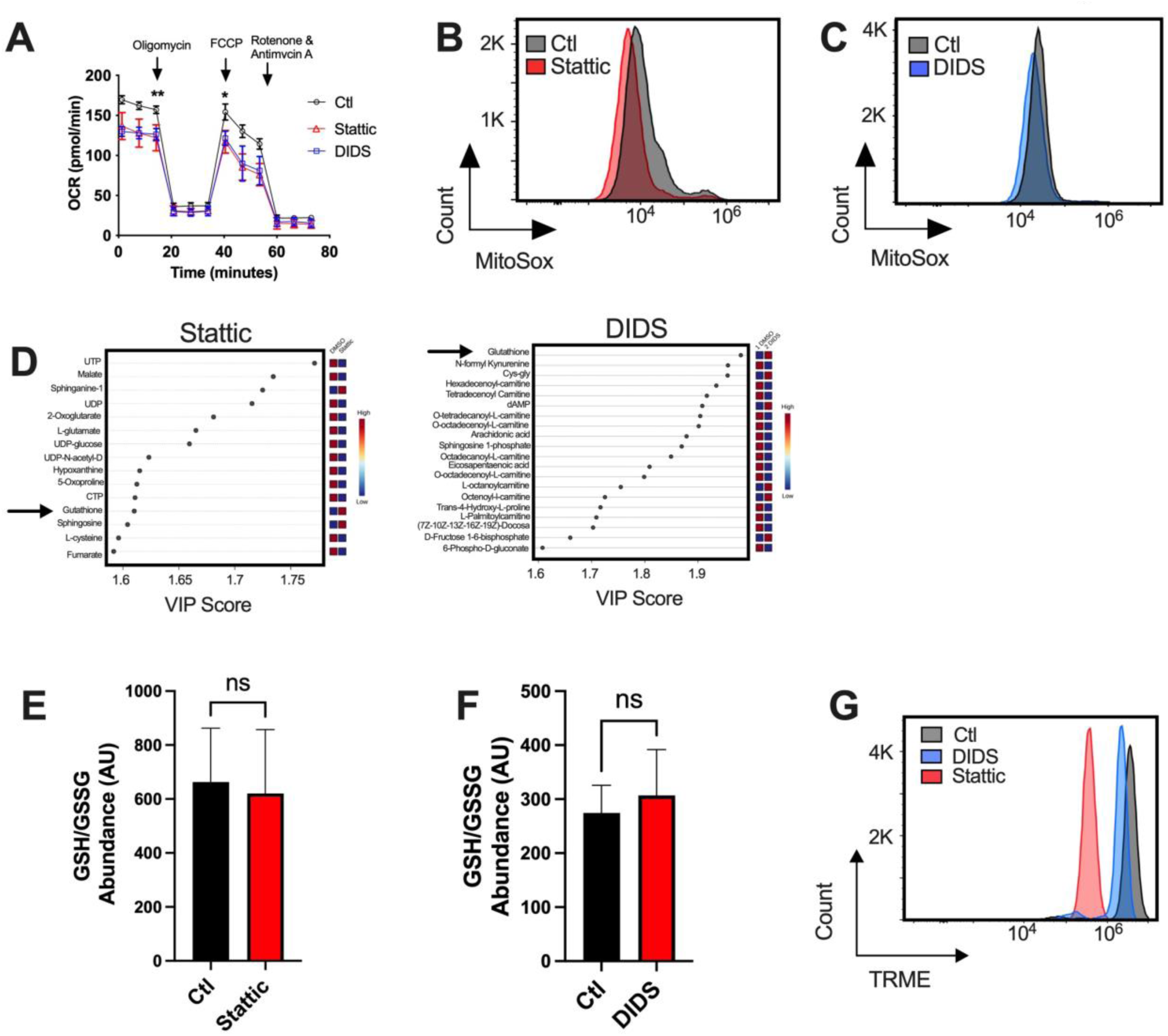
STAT3 and VDAC1 inhibition results in decreased OXPHOS and reduction in mitochondrial membrane potential. (A) Seahorse Mito Stress Test in MOLM-13 in the presence or absence of Stattic 5μM after 6 hours in culture. (B) Mitochondrial ROS as measured by flow cytometry in MOLM-13 cells treated with Stattic 5μM or vehicle control for 9 hours. (C) Mitochondrial ROS as measured by flow cytometry in MOLM-13 cells treated with DIDS 400μM or vehicle control for 9 hours. (D) VIP plots showing the top metabolites changed in global metabolomics of MOLM-13 cells treated with Stattic, DIDS compared to vehicle controls for 9 hours. (E) Glutathione (GSH) to glutathione disulfide (GSSG) ratio MOLM-13 cells treated with Stattic or DIDS for 9 hours compared to vehicle controls. (G) Mitochondrial membrane potential as measured by TMRE stain of MOLM-13 cells treated with Stattic or DIDS for 9 hours compared to vehicle control. Statistical analyses were performed using a Student’s t-test. P values are represented as follows: * p ≤ 0.05, ** p ≤ 0.01, *** p ≤ 0.001.

### STAT3 and VDAC1 inhibition leads to decreased in mitochondrial size

To determine whether the imbalance of calcium and OXPHOS result in mitochondrial dysfunction or changes in mitochondrial mass, we then studied how inhibition of STAT3 or VDAC1 affects the mitochondria, several hours after the decrease in mitochondrial calcium. Using electron microscopy, we found that pharmacologic inhibition of STAT3 with Stattic for 14 hours results in decreased mitochondrial size (**Figure 4A**). While Stattic-treated cells showed no significant difference in mitochondrial number per cell (**Figure 4B**), there was a significant decrease in mitochondrial size based on quantification of mitochondrial area (**Figure 4C**). Similarly, using mitotracker green stain, we determined that STAT3 inhibition with Stattic or siRNA knockdown resulted in decreased mitochondrial mass (**Figure 4D and Supplemental Figure 4)** when compared to controls, and VDAC1 inhibition showed similar results (**Figure 4E**). Taken together, these changes suggest that STAT3 of VDAC1 inhibition alters mitochondrial mass, likely through mitophagy.

**Figure 4.**
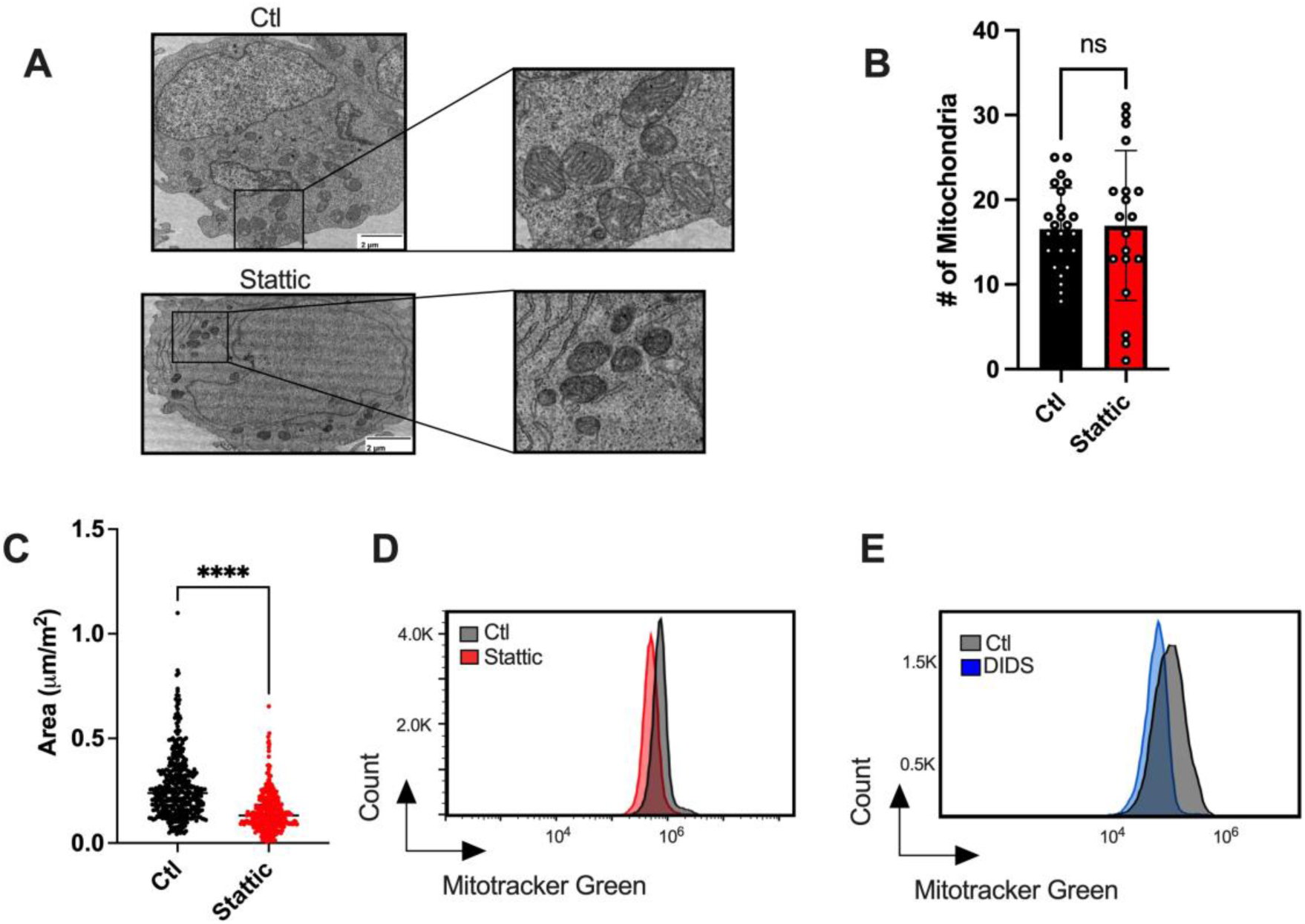
STAT3 and VDAC1 inhibition leads to decreased mitochondrial size. (A) Representative electron microscopy images of MOLM-13 cells treated with of Stattic 5μM for 14 hours or vehicle control. (B) Quantification of number of mitochondria imaged by electron microscopy in MOLM-13 cells treated with of Stattic 5μM for 14 hours or vehicle control. (C) Quantification of mitochondrial area of MOLM-13 cells treated with of Stattic 5μM for 14 hours or vehicle control. (D) Mitochondrial mass as measured by flow cytometry with mitotracker green stain in the presence of Stattic 5μM for 14 hours compared to vehicle control. (E) Mitochondrial mass as measured by flow cytometry with mitotracker green stain in the presence of DIDS 400μM for 14 hours. Statistical analyses were performed using a Student’s t-test. P values are represented as follows: * p ≤ 0.05, ** p ≤ 0.01, *** p ≤ 0.001.

### STAT3 and VDAC1 inhibition decreases viability and engraftment potential of leukemic cells

To determine the impact of STAT3 inhibition on leukemic cells *in vitro*, we cultured MOLM-13 cells with Stattic 5mM or vehicle control for 24 hours, followed by flow cytometry viability assays. As shown in **Figure 5A**, there was significant cell death of Stattic treated cells. Similarly, we saw significant cell death in MOLM-13 cells treated with DIDS for 24 hours compared to vehicle control (**Figure 5B**). Interestingly, culturing MOLM-13 cells with both Stattic and DIDS resulted in similar cell death, suggesting they are acting through a common pathway. We then sought to understand whether LSCs from primary AML samples would be sensitive to STAT3 inhibition. To do so, we used LSCs isolated from three different AML patient samples that were notably resistant to venetoclax. We then treated LSCs *in vitro* with Stattic 5mM or vehicle control for 16 hours, followed by viability assays. LSCs were independently treated with venetoclax as a positive control to ensure resistance. As shown in **Figure 5C**, we saw a significant decrease in viability with Stattic treatment compared to vehicle control. To further assess the effect of STAT3 and VDAC1 inhibition in leukemia stem cells, we treated three AML patient samples *ex-vivo* with Stattic 5mM or vehicle control for 16 hours, followed by xenotransplantation into NSG-S mice pre-treated with Busulfan. After 8-12 weeks, we found almost complete eradication of LSCs treated with Stattic, reflected by limited engraftment compared to vehicle controls in all three AML samples (**Figure 5D**). *Ex-vivo* VDAC1 inhibition with DIDS also resulted in a significant decrease in engraftment potential of two AML samples (**Figure 5E**). To determine whether STAT3 or VDAC1 inhibition results in a detrimental effect in normal hematopoietic stem cells (HSCs), we cultured CD34+ isolated cells from three adult bone marrow samples with Stattic or DIDS overnight followed by colony forming assays. As shown in **Figure 5F**, STAT3 inhibition results in decreased colony forming potential in HSCs, while VDAC1 inhibition results in increased colony forming potential suggesting a possible protective effect.

**Figure 5.**
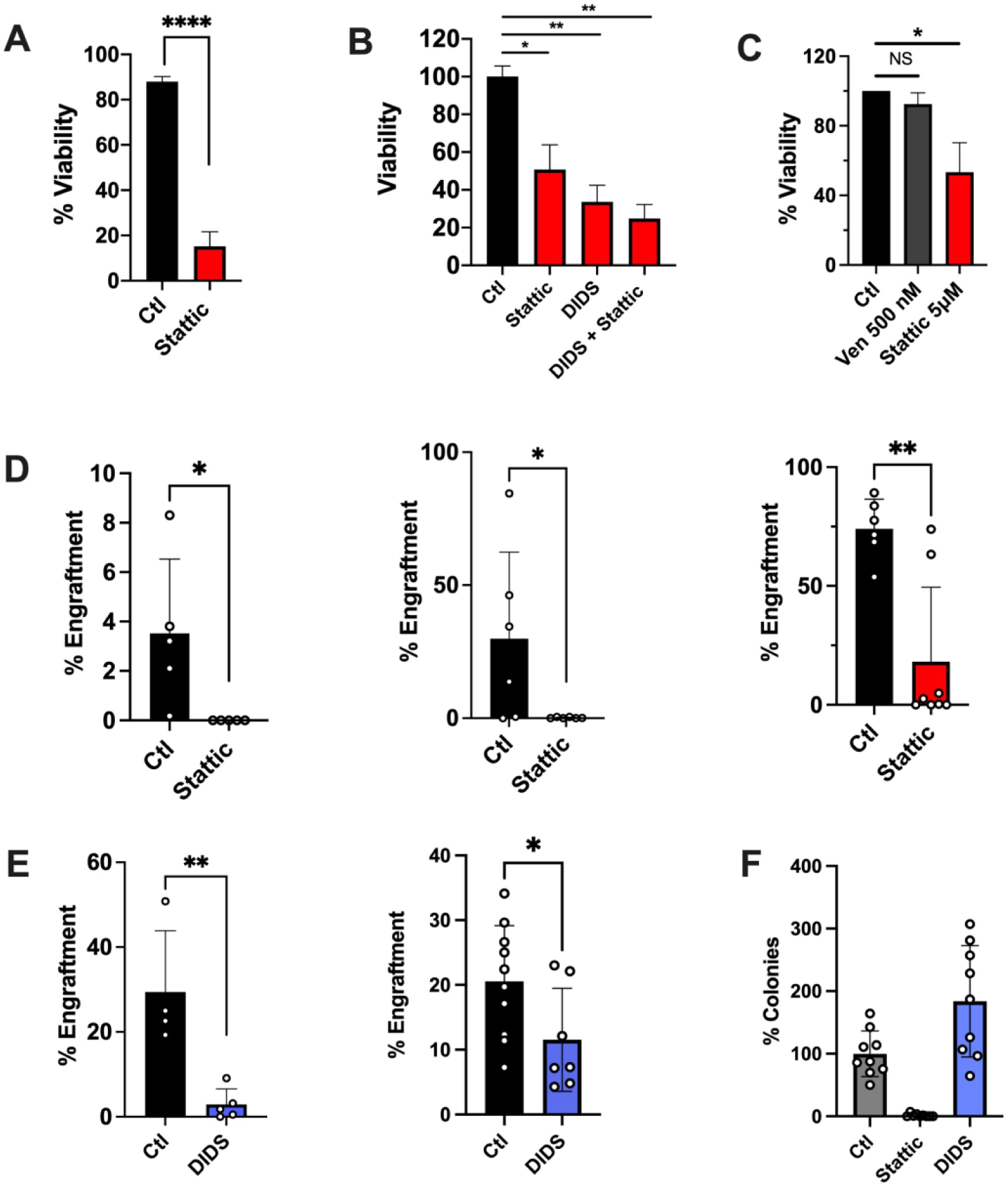
STAT3 and VDAC1 inhibition decreases viability of AML cells and it impairs leukemia stem cell function. (A) Viability as measured by flow cytometry of MOLM-13 cells treated with Stattic 5μM or vehicle control for 24 hours. (B) Viability as measured by flow cytometry of MOLM-13 cells treated with Stattic 5μM, DIDS 400μM, the combination or vehicle control for 24 hours. (C) Viability as measured by flow cytometry of ROS-low enriched leukemia stem cells isolated from AML patient samples and treated with venetoclax 500nM, Stattic 5μM or vehicle control for 24 hours. (D) Engraftment as measured by human CD45 positivity in NSG-S mice injected with bone marrow cells from three primary AML samples treated *ex-vivo* with Stattic 5μM or vehicle control for 16 hours. (E) Engraftment as measured by human CD45 positivity in NSG-S mice injected with bone marrow cells from two primary AML samples treated *ex-vivo* with DIDS 400μM or vehicle control for 16 hours. (F) Colony forming potential of CD34+ isolated cells from normal bone marrow samples and treated with Stattic 5μM, DIDS 400μM or vehicle control for 16 hours followed by methylcellulose plating.

## Discussion

STAT3 is a well-established transcription factor which is known to play important roles in cancer cell transformation and expansion^24,48^. In LSCs, STAT3 has been shown to regulate both MCL-1 expression^25^ and glutamine flux^26^, which are critical pathways for LSC survival. STAT3 phosphorylation at S727 has recognized importance in both the function of mitochondrial STAT3, and localization of STAT3 to the mitochondria^30^. While recent research has shown that mitochondrial STAT3 also plays a role in regulating ETC activity^30–32^ and mitochondrial calcium flux^34,35^, this function has not been studied in the context of myeloid malignancies, and the link between STAT3 and calcium regulation has not been well defined.

In this study, we confirm prior reports showing STAT3 in highly expressed in AML patient samples and cell lines, and that pSTAT3, in particular serine 727 phosphorylation, is associated with mitochondrial function of STAT3. We show that transcriptomic and pharmacologic inhibition of STAT3 results in impaired OXPHOS and decreased mitochondrial size. While the role of STAT3 in OXPHOS has been demonstrated given its regulation of *MYC* and downstream glutaminolysis^26^, we also discovered a novel role of mitochondrial STAT3 via its interaction with VDAC1. VDAC1 has a variety of mitochondrial roles, including regulating mitochondrial calcium^46^. In this study, we show that STAT3 inhibition leads to decreased VDAC1 in the mitochondrial cell fractions, followed by a decrease in mitochondrial calcium content (**Figure 6**). However, VDAC1’s role in apoptosis appears to be independent of STAT3 as the development of apoptosis does not appear to be affected by STAT3 inhibition. Further, VDAC1 inhibition effectively results in mitochondrial dysfunction and decreased mitochondrial mass, followed by cell death of AML cells, suggesting the STAT3-VDAC1 interaction is a critical component of the cell’s survival. Importantly, we show that STAT3 inhibition effectively kills LSCs, thereby impairing their engraftment potential. Similar to other studies^48^, we additionally demonstrate that STAT3 inhibition can be an effective modality to overcome venetoclax resistance, which aligns with prior work that has demonstrated the importance of mitochondrial calcium in venetoclax-resistant LSCs^19^. Interestingly, while inhibiting all functions of STAT3 through Stattic may be detrimental to HSCs, VDAC1 inhibition results in no decrease in HSC colony forming potential, suggesting targeting this specific pathway may be a potential good strategy in AML.

**Figure 6.**
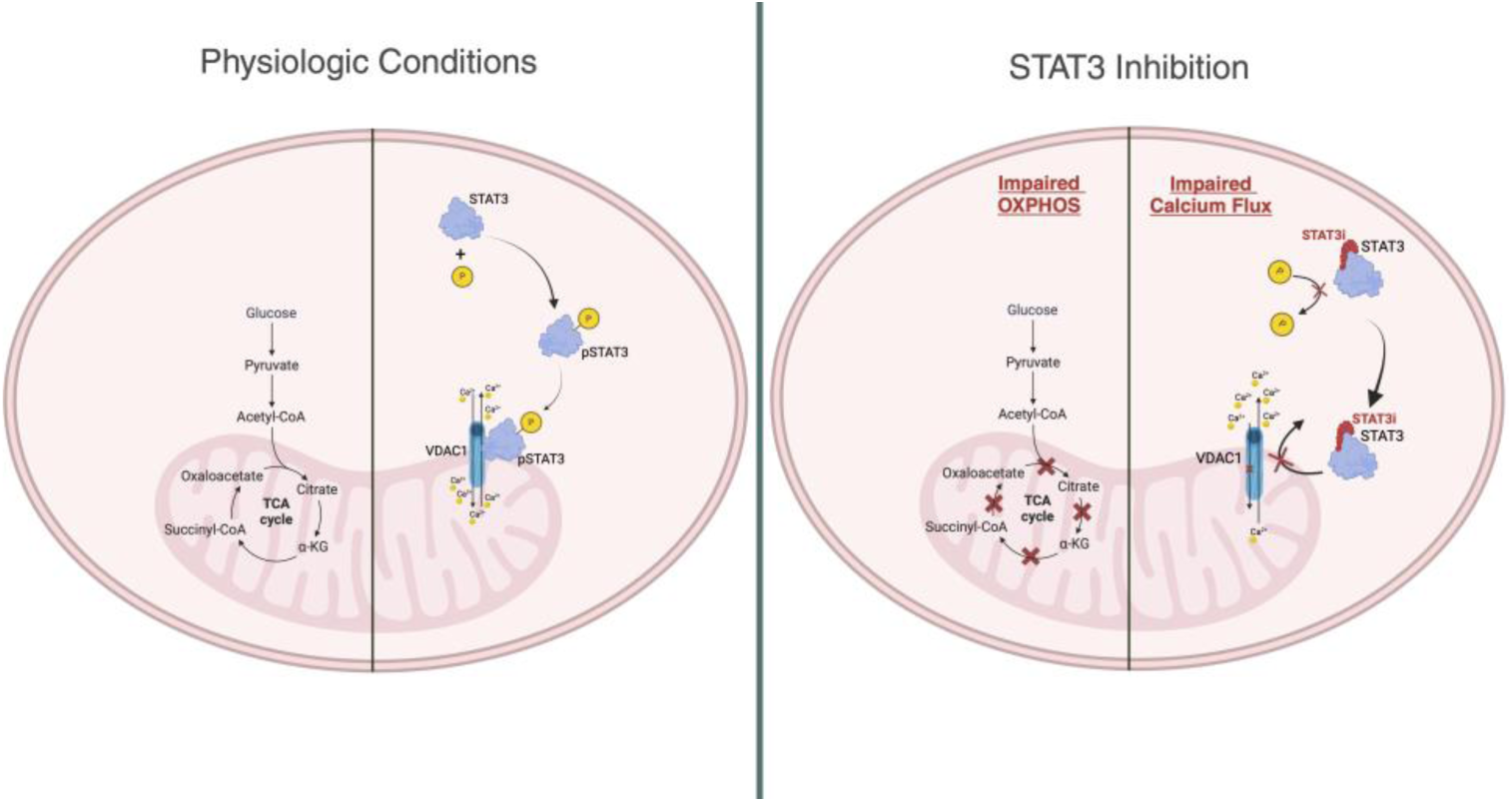
Model of STAT3 and VDAC1 Roles in AML cells. STAT3 is phosphorylated at S727 site and it localizes to the mitochondria, where it interacts with VDAC1 to regulate calcium intake into the mitochondria. Following STAT3 inhibition, this interaction is disrupted, leading to less mitochondrial calcium and mitochondrial dysfunction. Cartoon was created with BioRender.

In conclusion, we show STAT3 is highly expressed in AML cells, and inhibition of STAT3 results in decreased OXPHOS, decreased mitochondrial calcium and mitochondrial mass, decreased cell viability, and impaired engraftment potential. We additionally describe a novel role of STAT3 which interacts with VDAC1 in the mitochondria. These important functions of STAT3 represent potential therapeutic strategies in targeting AML, including LSCs. Given the promising therapeutic implications of targeting STAT3, inhibitors of this protein are currently being investigated in AML (NCT05986240) as well as other cancers (NCT03195699).

## Acknowledgements

We would like to thank all patients and their families who donated specimens to this research. We also thank the University of Colorado Mass Spectrometry Metabolomics and Proteomics Cores, and the Rocky Mountain Regional VA Flow Cytometry Core for their support. This work was supported by the Veteran’s Affairs CDA-2 (grant BX005603-01A1) (M.L.A.) and NHLBI grant 2R38HL 143511-05 (K.B.G.).

## Authorship Contributions

KG and MLA designed the research, KG, JB, RP, AI, GA, JR, AG, WS, and AD performed experiments and analyzed the data. KG and MLA wrote the manuscript with input from AEG, CS, AD, CM and DP

