## Supplement Figures for "The STAT3-VDAC1 Axis Modulates Mitochondrial Function and Plays a Critical Role in the Survival of Acute Myeloid Leukemia Cells"

**SUPPLEMENTAL MATERIALS**

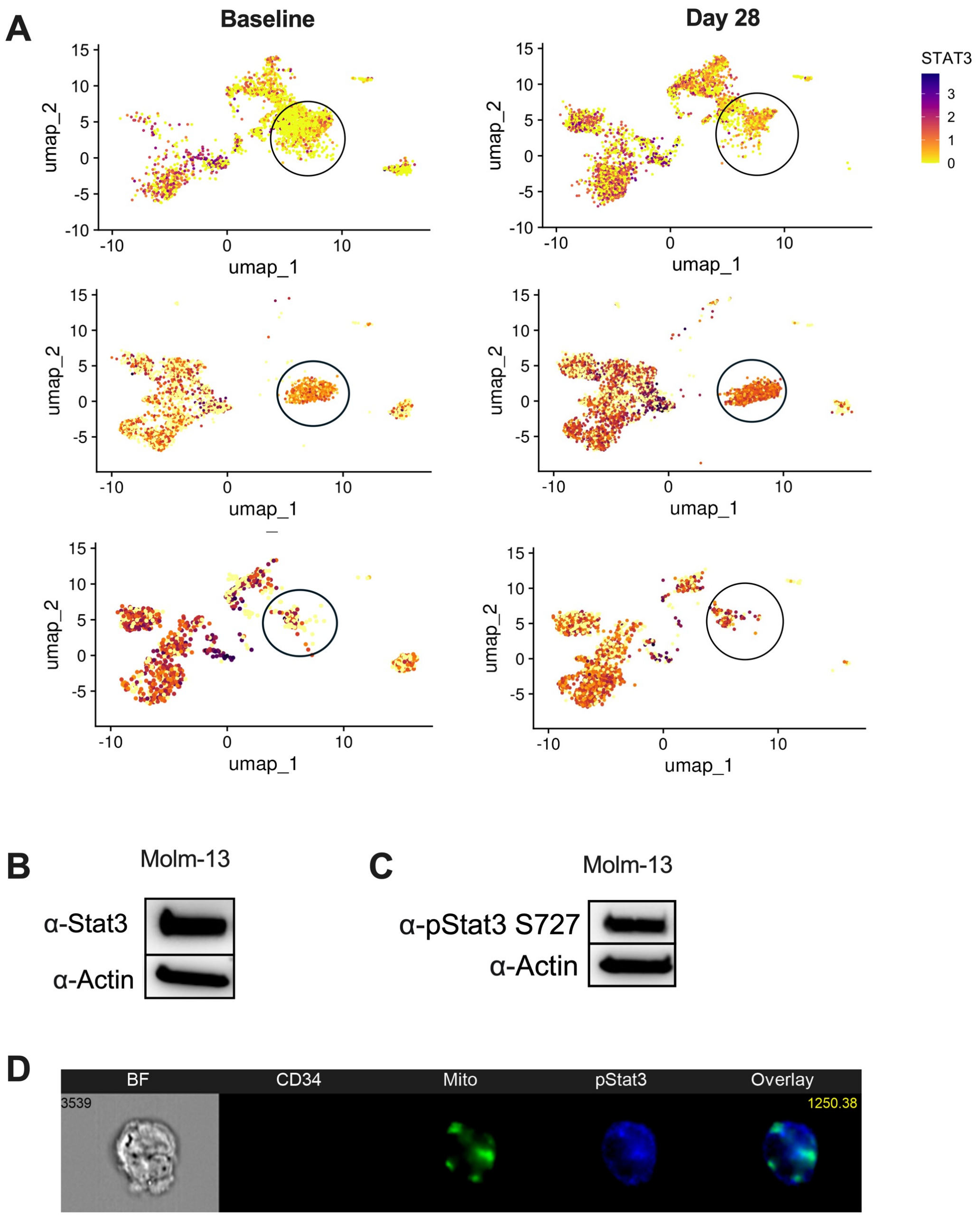


**Supplemental Figure 1**. **STAT3 is expressed in AML samples resistant to venetoclax and it localizes to the mitochondria of AML cells.** (A) STAT3 expression based on single cell RNA sequencing from 3 AML samples. Circled cluster shows leukemia stem cells. (B) Western blot showing STAT3 expression in Molm-13 cells. (C) Western blot showing phosphorylated STAT3 expression at the S272 site. (D) Image from a cell showing STAT3 localization to the mitochondria in Molm-13 cells.

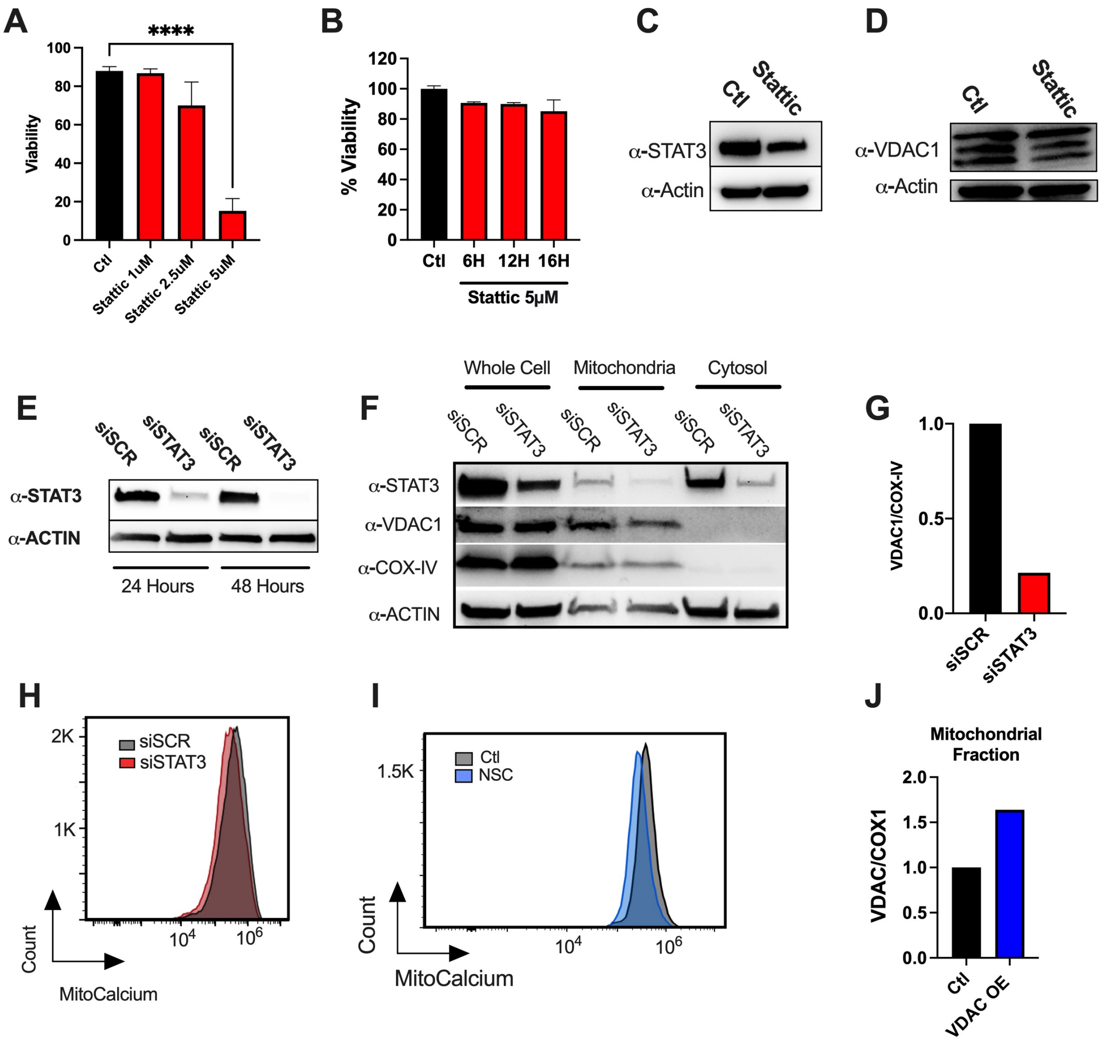


**Supplemental Figure 2. STAT3 regulates mitochondrial VDAC1 and affects calcium metabolism.** (A) Viability as measured by flow cytometry of Molm-13 cells treated with increasing doses of Stattic for 24 hours. (B) Viability as measured by flow cytometry of MOLM-13 cells treated with Stattic 5μM at various timepoints. (C) STAT3 protein expression by western blot in Molm-13 after treatment with Stattic 5μM for 9 hours or vehicle control. (D) VDAC1 protein expression by western blot in MOLM-13 cells treated with Stattic 5μM for 9 hours or vehicle control. (E) STAT3 protein expression by western blot in MOLM-13 cells treated siRNA against STAT3 or scrambled control for 24 and 48 hours. (F) Western blot showing protein expression of STAT3 and VDAC1 in whole cell, mitochondrial or cytosolic fractions of Molm-13 cells treated with siRNA against STAT3 or scrambled control for 48 hours. COX-IV and Actin antibodies serve as mitochondrial and cytosolic controls, respectively. (G) VDAC1/COX-IV quantification from western blot in (F). (H) Mitochondrial calcium as measured by flow cytometry of MOLM-13 cells treated with siSCR or siSTAT3 for 48 hours. (I) Mitochondrial calcium as measured by flow cytometry of MOLM-13 cells treated with the VDAC1 inhibitor NSC or vehicle control for 9 hours. (J) Quantified protein expression based on western blot of VDAC1 in mitochondrial fractions of MOLM-13 cells treated with VDAC1 overexpression vector or vector control. COX1 loading control expression was used for normalization.

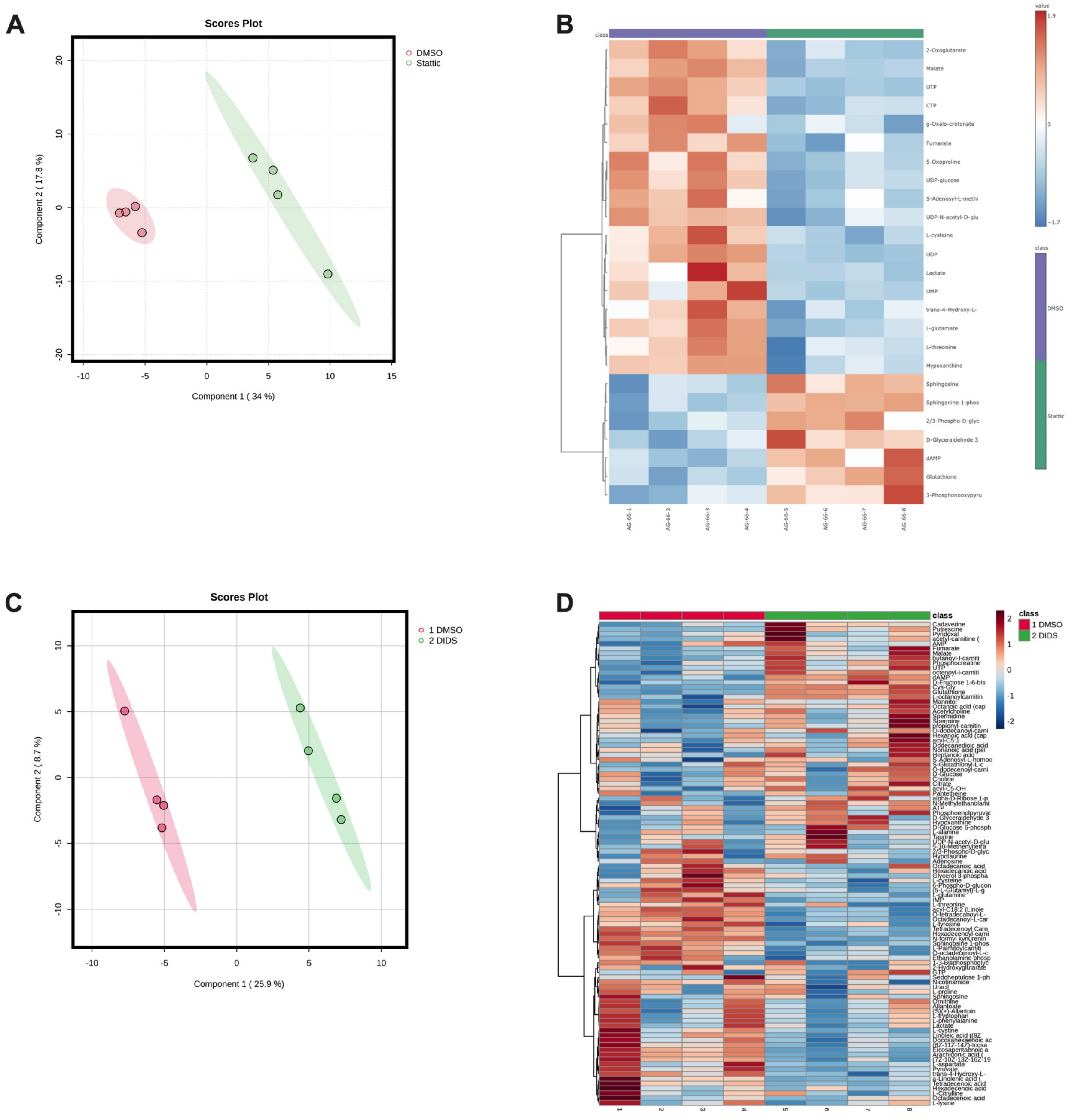


**Supplemental Figure 3. STAT3 and VDAC1 inhibition results in altered metabolic properties in AML cells**. (A) Scores plot from global metabolomics of MOLM-13 cells treated with 9 hours of Stattic 5μM or vehicle control. (B) Heatmap showing altered metabolites of MOLM-13 cells treated with 9 hours of Stattic 5μM or vehicle control. (C) Scores plot from global metabolomics of MOLM-13 cells treated with 9 hours of the VDAC1 inhibitor DIDS 400μM or vehicle control. (D) Heatmap showing altered metabolites of MOLM-13 cells treated with 9 hours of DIDS 400μM or vehicle control.

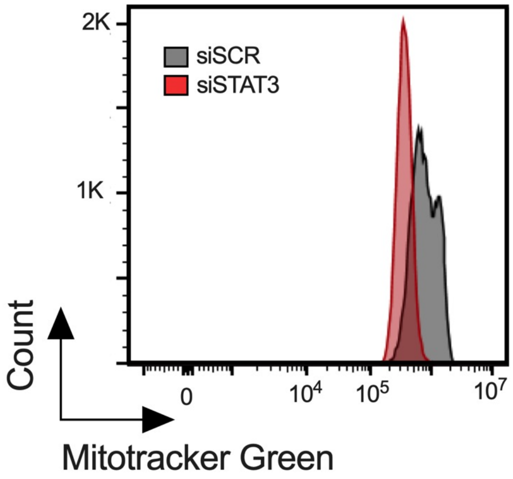


**Supplemental Figure 4**. **STAT3 inhibition leads to decreased mitochondrial size.** Flow cytometry showing mitotracker green stain of MOLM-13 cells treated with siSCR or siSTAT3 for 48 hours.
